# An integrated pipeline to estimate transcripts counts with single-cell resolution in *Caenorhabditis elegans*

**DOI:** 10.64898/2026.04.15.718387

**Authors:** Eva Sheardown, Yan To Ling, Servaas N. van der Burght, Heli Vaikkinen, Beth Gowing, Julie Ahringer, Fursham Hamid, QueeLim Ch’ng

## Abstract

Quantifying transcript abundance at single-cell resolution is important for understanding gene regulation in intact multicellular organisms. In *Caenorhabditis elegans*, RNA fluorescence in situ hybridization (RNA-FISH) has been widely used to visualize transcripts, but conventional single-molecule fluorescence in situ hybridization approaches can be limited by low signal-to-noise, poor labelling for short transcripts, and lack of streamlined quantitative analysis in identified cells. To overcome these issues and to simplify adoption, we designed an integrated experimental and computational pipeline for quantitative transcript analysis in *C. elegans* that exploits the increased sensitivity of Hybridisation Chain Reaction RNA-FISH and the new automated spot-detection software. This approach enables multiplexed detection of transcripts with single-cell resolution that agree with known spatial expression patterns in embryos, larvae, and adults. Applying this pipeline to the short insulin-like peptide *ins-4* in specific sensory neuron pairs in mutant backgrounds revealed quantitative changes in gene expression. This pipeline provides an accessible method to uncover spatial patterns of transcript localization and estimates of transcript counts in anatomically intact *C. elegans* to support mechanistic studies of gene regulation.

**Summary:** Measuring gene activity is a key to genetics research. Here, we combine cutting-edge tools for visualizing and counting mRNAs for a given gene with single-cell resolution in the nematode *C. elegans*. The resulting pipeline works for any developmental stage, has high signal-to-noise, can analyze multiple genes simultaneously, and is semi-automated. We demonstrate that approach can reveal differential effects of a mutation on gene expression in different cells. We hope that this inexpensive and user-friendly solution will increase adoption of improved techniques in the *C. elegans* community, while providing generalizable analysis approaches to a wider scientific audience.

## Introduction

Gene expression plays key roles throughout biology, controlling diverse processes from cell fates to organismal states. Transcription is major point of regulation during gene expression, with transcript abundance being one of the most common readouts of gene activity (Cramer 2019; Pope and Medzhitov 2018). Determining the spatial abundance of different transcripts in whole-mount samples has been especially valuable in studies of multicellular organisms where gene activity can be ascribed to specific cell and tissue types (Calarco et al. 2025).

RNA fluorescence in situ hybridization (RNA-FISH) has emerged as a key method to visualize individual transcripts in anatomically intact samples (Femino et al. 1988; Lee et al. 2017; Raj et al. 2008). The general approach of RNA-FISH is to label single transcripts with many fluorescent molecules that together produce a detectable signal visible under fluorescence microscopy as a diffraction-limited spot. This technique has myriad applications in investigating gene regulation, RNA localization, transcriptional noise, inter-cellular heterogeneity, and other RNA-related processes (Baudrimont et al. 2017; Buxbaum et al. 2015; Duncan and Rosa 2018; Ercan et al. 2009; Omerzu et al. 2019; Paré et al. 2009; Trimmer et al. 2023). It has also provided a rapid way to validate results from transcriptomic experiments (Cheng et al. 2023). Over the last decade, advances that enhance sensitivity, specificity, multiplexing, and image analysis of RNA-FISH have simplified the quantification of transcript abundance. It is now feasible to count multiple transcripts with near single-molecule accuracy within individual cells in whole-mount samples (Choi et al. 2010; Shah et al. 2016; Zhao et al. 2023). This ability facilitates the transition from qualitative to quantitative data necessary for mechanistic, systems understanding.

Transparent preparations are particularly amenable to RNA-FISH labelling, dispensing with the need for steps such as sectioning or clearing prior to imaging. Transparency is one of the many advantages of nematode *C. elegans*, a major model system used extensively in genetic studies of gene expression and regulation, from single-gene analysis to transcriptomics. *C. elegans* also possesses the advantage of a fully defined and stereotyped anatomy, where every somatic cell is identified and where individuals possess the identical set of cells, enabling labelled transcripts to be associated with cells of specific identities (Cao et al. 2017; Packer et al. 2019; Sulston et al. 1983).

One of the earliest single molecule RNA-FISH techniques was developed in *C. elegans* (Raj et al. 2008). Since then, smFISH has been used successfully in many *C. elegans* studies and remains the most widely used RNA-FISH techniques in *C. elegans* (Ji and van Oudenaarden 2012; Korzelius et al. 2011; Lee et al. 2019; Parker et al. 2021; Saffer et al. 2011; Winkenbach et al. 2022; Yu et al. 2020). Nonetheless, these smFISH implementations present three common technical limitations. First, the signal-to-noise is not as high as newer methods, which restricts signal detection under high background fluorescence. This issue is particularly vexing in *C. elegans* larvae and adults, which tend to exhibit more autofluorescence. Second, smFISH relies on hybridizing many fluorescent oligonucleotide probes across a single transcript (Raj et al. 2008). Short transcripts cannot accommodate as many unique probes, which makes it challenging to visualize them. Third, some studies rely on total fluorescence within a region of interest (ROI) as a proxy for transcript abundance. This approach loses the ability to count transcripts and can produce noisier readouts as individual transcripts differ in fluorescence levels due to background and variable signal amplification (Battich et al. 2015; Raj et al. 2006; Raj et al. 2008)

These limitations are surmountable through newer RNA-FISH and spot-detection methods. Hybridization Chain Reaction (HCR) is a well-established RNA-FISH technique with greater signal amplification through self-assembly of many fluorescent labels onto probes complementary to different parts of a transcript (Choi et al. 2010; Choi et al. 2014; Dirks and Pierce 2004; Shah et al. 2016; Yang et al. 2026). HCR has been well established across diverse sample types, including *C. elegans,* allowing for imaging of RNA molecules at sub-cellular resolution (Choi et al. 2016) and the use of split-initiator probe pairs developed in HCR3.0 created better signal-to-noise allowing for single molecule counting (Choi et al. 2018; Shah et al. 2016). HCR requires fewer distinct binding sites, making it suitable for labelling shorter transcripts. Additionally, software tools such as RS-FISH have improved the ability to identify fluorescent spots corresponding to labelled transcripts (Bahry et al. 2022).

HCR3.0’s ability to label single molecules of mRNA with high efficiency, specificity, and signal-to-noise is well-validated, as is the precision and effectiveness of RS-FISH in spot detection. However, these techniques are not widely adopted in *C. elegans* studies, likely because of costs, the need for optimization, and the disparate experimental and computational steps required. We were therefore motivated to address this practical gap. To facilitate adoption, we developed an experimental and analysis pipeline that adapts whole mount preparation across multiple *C. elegans* developmental stages, detects short transcripts with reduced probe-pair numbers, and links HCR3.0 chemistry and imaging to RS-FISH spot detection to produce transcript counts in individual identified cells in *C. elegans*.

## Results

### An integrated pipeline for quantitative HCR analysis in *C. elegans* embryos, larvae, and adults

HCR provides improved signal-to-noise compared to conventional smFISH. However, one barrier in applying quantitative HCR analysis to current *C. elegans* research is the substantial effort and range of expertise needed to optimize fixation and labelling, while developing computational approaches to identify fluorescent spots and quantifying them. Building on previous whole-mount *C. elegans* HCR work (Choi et al. 2016) and the quantitative abilities of HCR3.0 (Choi et al. 2018) we have combined these steps in a single pipeline (Figure 1) to reduce barriers to HCR adoption. These steps are detailed in the materials methods and the accompanying protocol.

**Figure 1.**
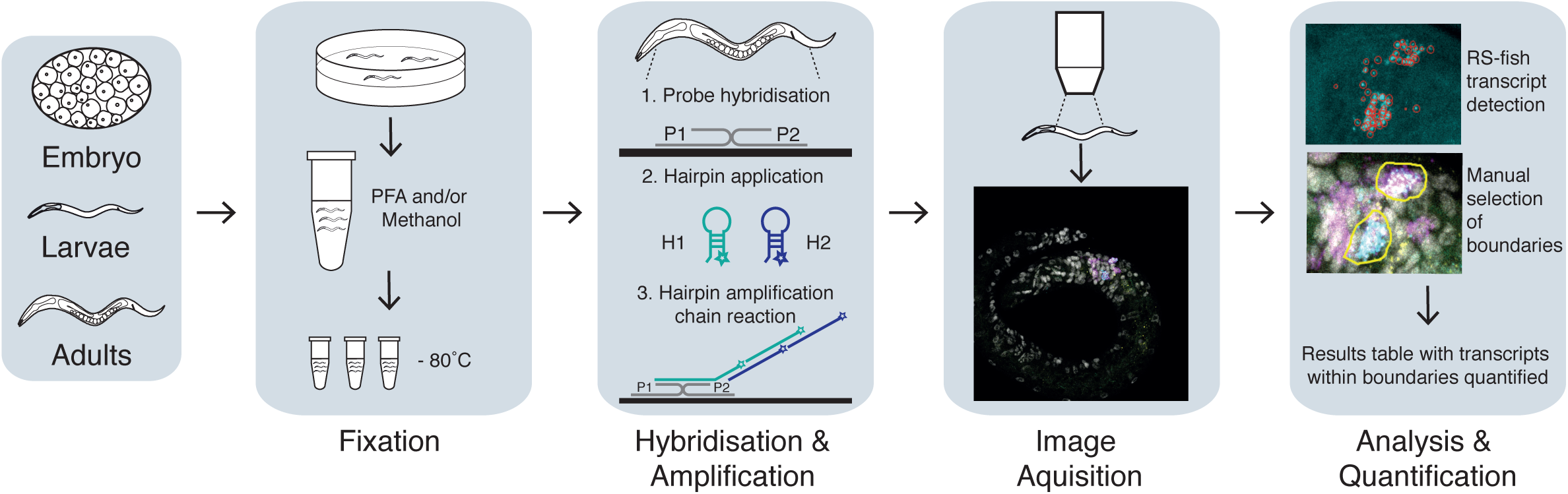
HCR Pipeline. A scheme of the overall pipeline. The inputs to this pipeline are embryos, larvae, or adults, which are frozen, fixed, labelled, and imaged. Spot detection and selection of a region of interest (which may correspond to single cells or tissues) are then applied to the image stacks. The outputs are the number of fluorescent spots corresponding to individual transcripts within the region of interest.

In this pipeline, embryos, larvae, or adults are harvested, frozen at -80°C with fixative, thawed, fixed, then hybridized to probes followed by signal amplification via HCR3.0 (see Methods). *C. elegans* embryos, larvae, and adults are processed differently due to anatomical differences such as size and the presence of an eggshell in embryos versus a cuticle in larvae and adults. Labelled samples can be stored for ∼4 weeks at 4°C in the dark.

Next, we acquire image stacks using fluorescence confocal microscopy, to maximize the signal from diffraction-limited fluorescent spots corresponding to individual mRNA molecules while minimizing out-of-focus background. To visualize multiple transcripts in the same individual, we use probes labelled with distinct fluorophores matched against the illumination wavelengths of the confocal microscope. We recommend using high numerical aperture objectives to maximise the ability to resolve fluorescent spots. Detailed methods for optimising imaging parameters are beyond the scope of this paper, but we recommend Waters (Waters 2009) as a comprehensive and practical guide.

Once the images are acquired, we use RS-FISH (Bahry et al. 2022) to identify fluorescent spots consistent with labelled transcripts. We then used FIJI (Schindelin et al. 2012) to select regions of interest (ROIs) to corresponding to specific cells. Spots within the ROI are counted by a MATLAB script to estimate the number of transcripts and their coordinates within a given cell. The output of this pipeline is the estimated number of transcripts and intensities within each cell (or a selected region of interest).

### Low-cost probe and amplifier design

HCR3.0 relies on multiple initiator DNA oligonucleotides with complementary sequences that enable them to bind to a target transcript (Choi et al. 2018). After binding, these oligonucleotides initiate an isothermal hybridization chain reaction that assembles multiple fluorescently labelled amplifier oligonucleotides, adding up to a detectable fluorescent signal. Using 15 or more probe pairs per transcript is recommended, although we have successfully visualized transcripts with as few as 4 probe pairs, allowing quantification of short transcripts for which larger probe sets are not feasible.

Molecular Instruments supplies kits for the entire HCR process. To save costs, it is possible to prepare buffers as well as design initiator probes and amplifier oligonucleotides separately. These pools of initiator probes and amplifier oligonucleotides can be ordered from any custom oligonucleotide synthesis company at a substantially lower cost.

Specific probe pairs complementary to a given target RNA sequence and B isoform are generated using a custom HCR probe designer (Suppermpool et al. 2025). Amplifier hairpins are designed using published sequences for the B isoforms (Choi et al. 2014) and selecting an Alexa Fluor of choice that matches the fluorescence channels on the microscope used.

### Multiplexed transcript visualization at single-cell resolution

To demonstrate the utility of this pipeline, we designed probes to quantify the abundance of multiple transcripts in embryos, second larval (L2) stage animals, and adults in whole-mount preparations. We compared formaldehyde with or without methanol fixation methods and found that including a methanol fixation step in embryos **|** nuclear morphology in methanol-fixed samples and collapsed nuclei in formaldehyde-fixed samples (Supplemental Figure 1). In L2 larval samples, permeability and detection rates were high (∼90%) with only formaldehyde fixation (Supplemental figure 2), however in adults yielded substantially higher detection rates with and additional a methanol fixation step (Supplemental Figure 3),

To illustrate multiplexing, we visualized the expression of the very early transcript *pes-10* (Seydoux and Fire 1994), the *Notch* target *ref-1* (Cole et al. 2024; Neves and Priess 2005), and the GATA factor *end-1* (Raj et al. 2010; Zhu et al. 1997) in early embryos. The HCR results showed the expected expression patterns at the 26-cell stage: strong *ref-1* expression in the D blastomere and the AB-lineage cells ABaraa and ABarap (and low expression in other somatic cells), *end-1* in the two E-lineage cells Ea and Ep, and *pes-10* in all somatic cells (Figure 2a-d).

**Figure 2.**
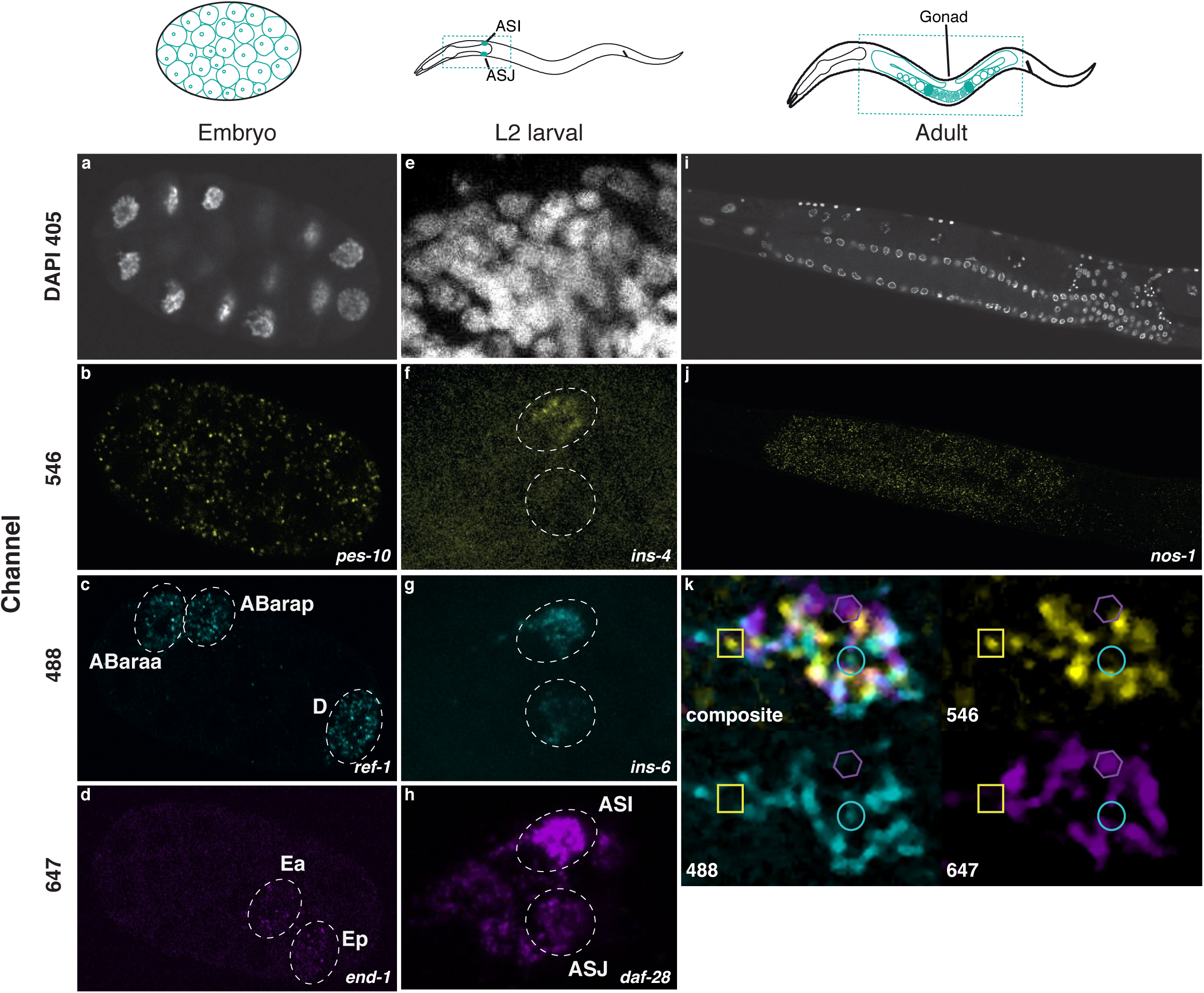
Multiplex labeling of individual transcripts in embryos, larvae, and adults. (a-d) a single slice image of a 26-cell embryo labelled with DAPI and probes for *pes-10*, *ref-1*, and *end-1*. The D blastomere, the AB-lineage cells ABaraa and ABarap and the two E-lineage cells Ea and Ep are indicated. (e-h). A representative maximum intensity projection of the right side of wild-type L2 larvae, labelled with DAPI and probes for *ins-4*, *ins-6*, and *daf-28* transcripts. ASI and ASJ sensory neurons are indicated in (h). (i-j) A single slice image of an adult worm labelled with DAPI and probes for *nos-1* localized within the germline syncytium. (k) A composite image merging multiple fluorescence channels showing minimal bleed-through. Individual channels also shown with channel specific transcripts indicated.

To apply our HCR pipeline to L2 larvae, we selected three insulin-like peptides (ILPs) for analysis: *ins-4*, *ins-6*, and *daf-*28 (Chen and Baugh 2014; Cornils et al. 2011; Fernandes de Abreu et al. 2014; Hung et al. 2014; Li et al. 2003). These ILPs are encoded by short mRNAs, which have limited their analysis using other RNA-FISH approaches. These ILPs are ideal for demonstrating the utility of this HCR pipeline for several reasons. First, their sites of expression have been previously characterized (Cornils et al. 2011; Hung et al. 2014; Li et al. 2003; Taylor et al. 2021; Wu et al. 2019), thus establishing clear positive controls. Second, their cell-specific expression emphasizes the spatial specificity of HCR labelling and minimal background. Finally, their overlapping expression patterns in the same neurons demonstrate minimal bleed-through between different fluorescence channels.

*ins-4* is primarily expressed in the ASI sensory neurons with modest expression in the ASJ sensory neurons (Hung et al. 2014; Taylor et al. 2021; Wu et al. 2019) whereas *ins-6* and *daf-28* are both expressed robustly in both ASI and ASJ sensory neurons (and in additional neurons) (Cornils et al. 2011; Hung et al. 2014; Li et al. 2003; Taylor et al. 2021). Both ASI and ASJ neurons occur as left-right symmetric pairs in the worm.

Figure 2e-h shows confocal images of an individual L2 larvae simultaneously labelled with probes against *ins-4*, *ins-6*, and *daf-28*. Spots corresponding to these transcripts are clustered around DAPI-stained nuclei corresponding to the ASI and ASJ neurons. The locations of these spots were consistent with the established expression patterns for *ins-4*, *ins-6*, and *daf-28*. HCR reveals *ins-4* transcripts in ASI with low levels in ASJ, high levels of *ins-6* transcripts in both ASI and ASJ, and *daf-28* transcripts in ASI and ASJ as well as additional head neurons.

In adults, we used HCR to visualize *nos-1*, a Nanos homolog expressed in the germline (Subramaniam and Seydoux 1999). Figure 2i-j shows *nos-1* transcripts in methanol-fixed adults, localized within the germline syncytium.

### Specific labelling of distinct transcripts

These results point to the specificity and multiplex capabilities of HCR. In embryos, only specific blastomeres are labelled with the relevant probes. In L2 larvae, there is relatively little staining outside the nervous system (e.g., skin) where these 3 ILPs are not expressed. These distinct spatial patterns also indicate good spectral separation among the probe sets for each transcript, which permits multiplexed analysis of different transcripts. We also have minimal bleed through between channels (Figure 2k), emphasizing specificity of labelling.

### Automated detection of labelled transcripts in single cells

To identify labelled transcripts, we integrated spot detection in our pipeline (Figure 4). Here, we used the well-established RS-FISH software to identify fluorescent spots corresponding to individual transcripts (Bahry et al. 2022). RS-FISH employs the Radial Symmetry (RS) algorithm to reliably detect and localize fluorescent spots with subpixel accuracy. The method is implemented as an ImageJ/Fiji plug-in, ensuring accessibility to the broader scientific community. It supports spot detection in three-dimensional images and accounts for anisotropy without the need for prior image rescaling. Furthermore, RS-FISH integrates Random Sample Consensus (RANSAC) for robust outlier removal, which enables accurate discrimination of spatially proximal spots.

Staining nuclei with DAPI allowed us to identify ASI and ASJ neurons in our L2 larvae image stacks and outline them using FIJI (Schindelin et al. 2012) to designate cell-specific ROIs. Using a custom MATLAB script, we identified the fluorescent spots detected by RS-FISH within each ROI. This analysis thus produces estimated transcript counts for each cell.

### Quantitative analysis of gene regulation

Next, we applied this pipeline to quantitative studies. Here, we exploited HCR to compare the cell-specific abundance of *ins-4* transcripts in ASI and ASJ neurons between wild type and mutant animals (Figure 3) (Barstead et al. 2012; Cornils et al. 2011; Fernandes de Abreu et al. 2014; Hung et al. 2014) in the L2 larval stage, where *ins-4* mRNA abundance has not been previously quantified. This analysis illustrates the application of the pipeline to compare short, low abundance transcript estimates.

**Figure 3.**
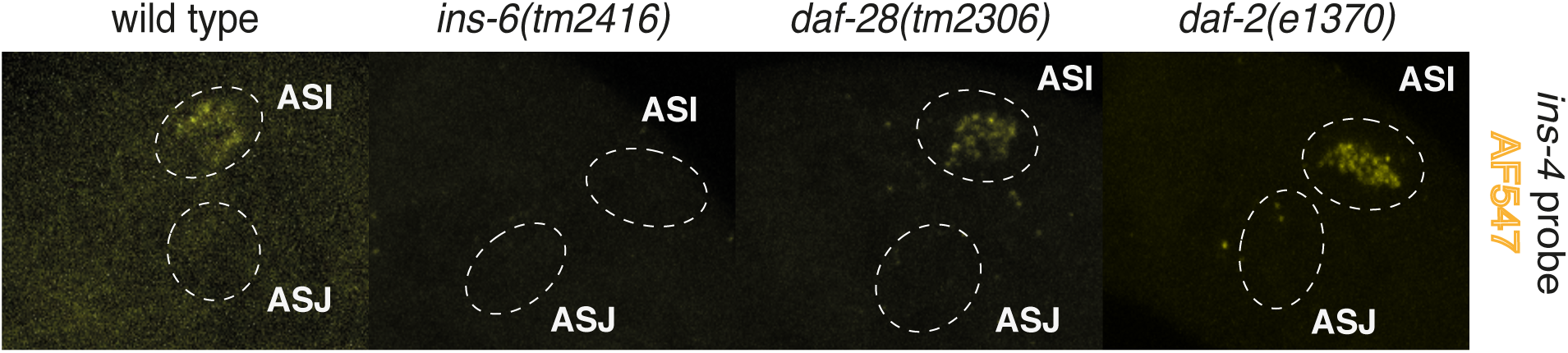
Transcript labelling in wild-type animals and deletion mutants. Representative maximum intensity projection images of the right side of wild-type, *ins-6(tm2416)*, *daf-28(tm2308)*, and *daf-2(e1370)* L2 larvae labelled with probes for *ins-4* transcripts. ASI and ASJ sensory neurons are circled.

**Figure 4.**
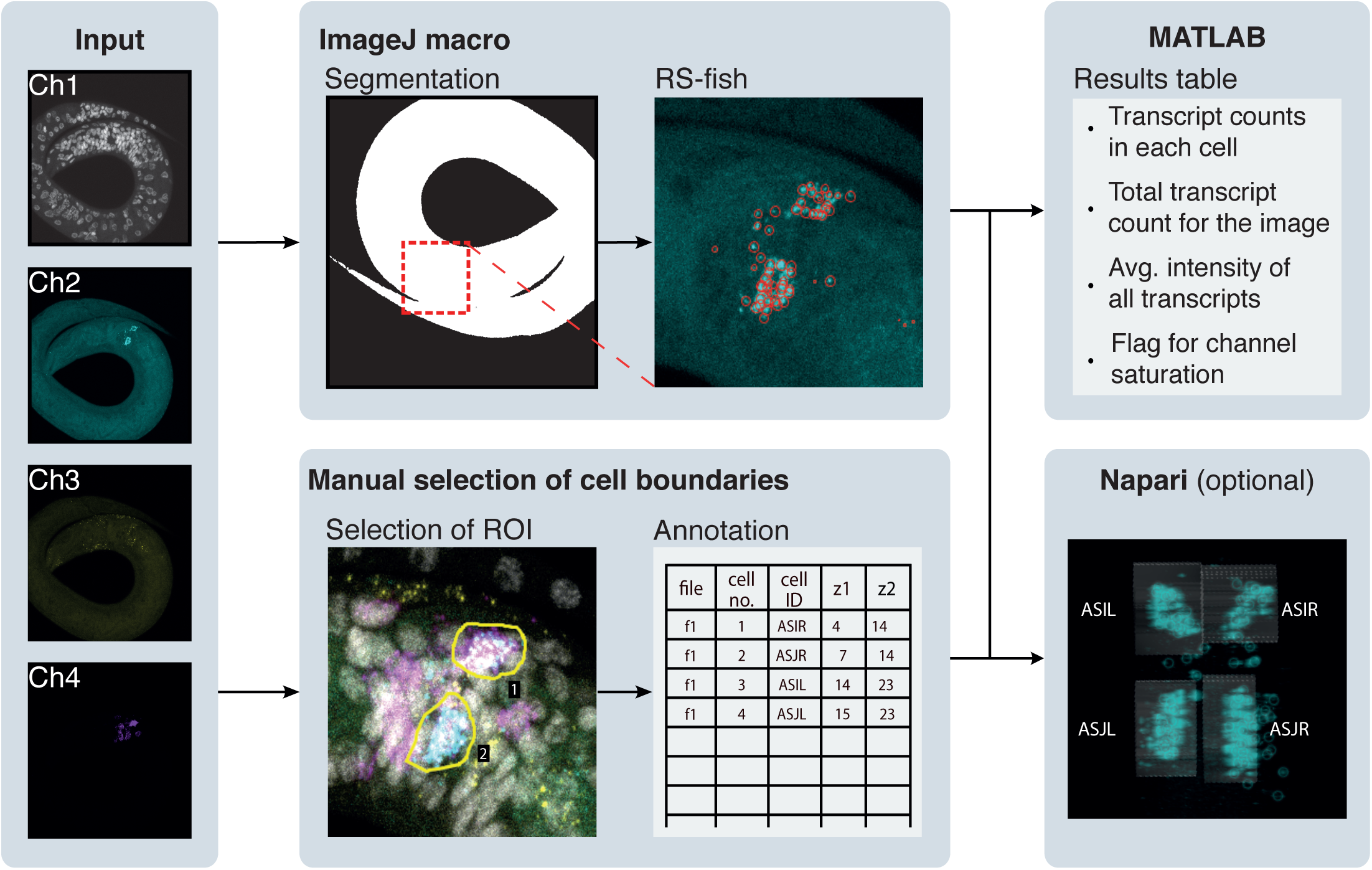
Image analysis to detect individual transcripts. Details of the image analysis pipeline to count individual transcripts. A 3D image was taken on a confocal microscope with the focal planes imaged determined using DAPI to find the dorsal and ventral points of the worm, ensuring the entire worm was captured. The ROI are then defined using ImageJ, and spot detection is performed with RS-FISH using the MATLAB script to select spots corresponding to transcripts within the ROI. A graphical user interface in Napari is included as an option for 3D visualization of the spots detected by RS-FISH and the volume selected for each cell.

We analysed the same image datasets using two quantitative methods: corrected mean fluorescence intensity within the neuronal ROI (Supplemental Figure 4) and RS-FISH-based transcript counting (Figure 5).

**Figure 5.**
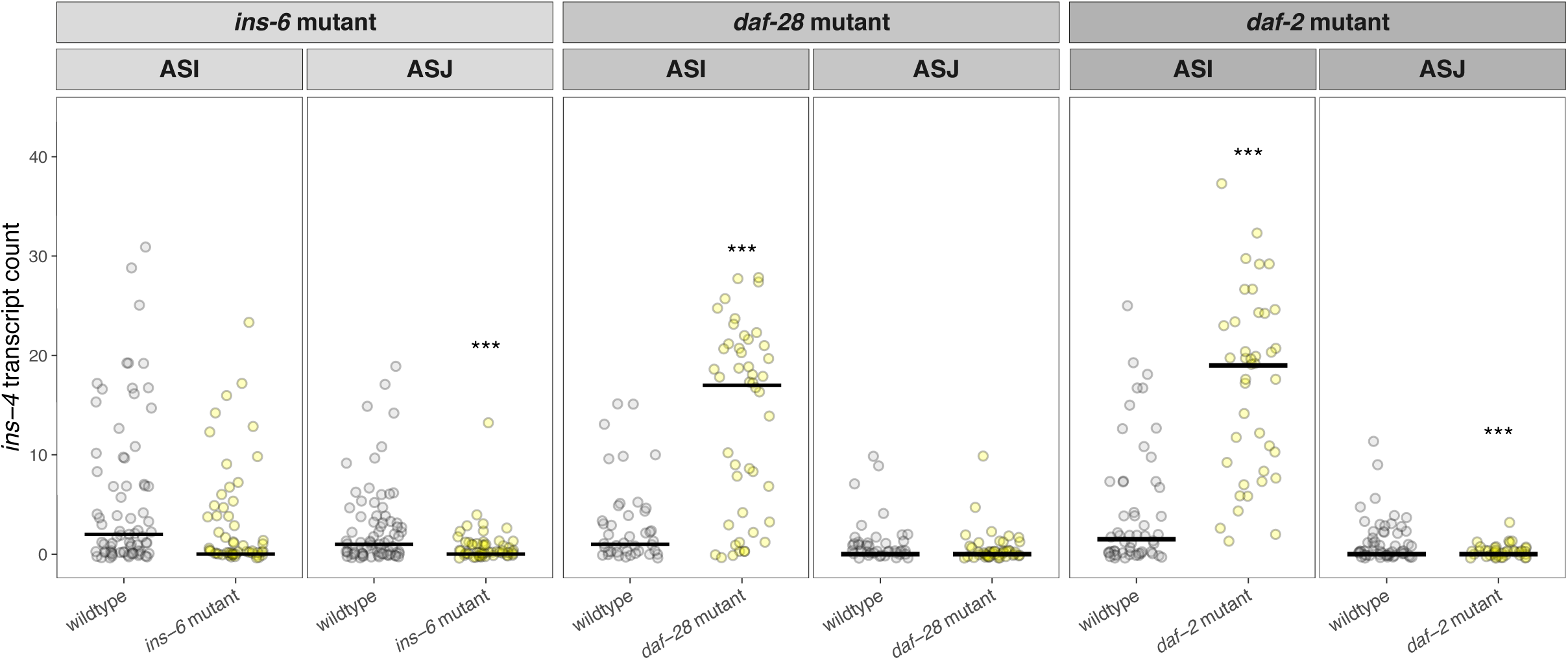
Transcript quantification in wild-type and deletion mutants. *ins-4* transcript quantification in individual ASI and ASJ neurons from wildtype and mutant L2 larvae. Columns show the mutant background: *ins-6(tm2416)*, *daf-28(tm2308)* and *daf-2(e1370)*. Grey and yellow points indicate wildtype and mutant genotype respectively. Each point represents the transcript count from a left and right neuron pair; horizontal black bars indicate the median. Asterisks indicate statistically significant differences between wildtype and mutant animals (*** P < 0.001). Further details in Supplemental Table 2.

Corrected mean fluorescence intensities are commonly used in quantifying expression levels of fluorescent molecules in transcriptional reporters and have been applied to RNA-FISH images. To quantify the corrected mean fluorescence intensity, we subtracted the mean background fluorescence from the mean fluorescence intensity measured within each neuronal ROI. This approach captures transcript-associated fluorescence within the cell but has several shortcomings (see Discussion).

HCR allows one to quantify transcript abundance by counting the number of fluorescent spots (Choi et al. 2018; Shah et al. 2016). In this pipeline, we can count RS-FISH detected puncta assigned to each cell to estimate of the number of transcript molecules present within a cell to near single molecule level. This approach is less sensitive to variations in fluorescence intensity between individual puncta.

Transcript abundance and corrected mean fluorescence generally identified similar changes in *ins-4* expression (Supplemental Figure 4 and Figure 5), although the magnitude and statistical significance of some effects differed. Generally, analyses based on counting RS-FISH detected puncta produced clearer separation between genotypes and identified a greater number of statistically significant differences than corrected mean fluorescence measurements alone. We further examined the sources of variance in these quantification approaches (see Methods) and found that the corrected mean fluorescence intensities were more sensitive to batch effects (Supplementary table 1). These findings suggest that counting transcript-associated puncta reduces measurement noise and improves robustness. We thus selected this approach as the primary quantitative readout.

In our transcript quantification, *ins-4* expression was significantly reduced in the ASJ but not ASI in *ins-6(tm2416)* mutants. Conversely, *ins-4* expression was increased in *daf-28(tm2308)* mutants compared to wildtype in ASI but not ASJ.

DAF-2 is the *C. elegans* ortholog of the mammalian insulin receptor (Kimura et al. 1997). In *daf-2(e1370)* mutants that reduce DAF-2 activity, *ins-4* expression was increased compared to wildtype in the ASI but decreased compared to wildtype in the ASJ. This result indicates that loss of *daf-2* produces distinct phenotypes in different neurons rather than a uniform response. This quantitative analysis also suggests that ASI and ASJ neurons operate independently and are affected differently by the same mutation.

## Discussion

Gene expression analysis is an essential part of a geneticist’s toolbox. Since the initial development of smFISH as a readout for gene expression, improvements in labelling (HCR3.0) and image processing techniques have enhanced the detection of single transcript molecules. Here we developed a pipeline that integrates HCR3.0 and RS-FISH to simplify the process of counting transcripts with single-cell resolution. We illustrated this pipeline’s multiplexed capability, specificity, quantitative output, and ability to visualize short transcripts. These results suggest several broad applications for this pipeline: identifying spatial patterns of gene expression, quantifying transcript abundance in identified cells across different genotypes, and simultaneously analyzing multiple transcripts to reveal their spatial and quantitative relationships.

Small genes with short transcripts such the ILPs lack favorable sites for the large number of probes required in smFISH. The ability to detect transcripts with only a few probes hybridizing presents opportunity to detect shorter transcripts that were not so easily detectible with other smFISH methods. Thus, our pipeline may be applicable to analysis of small coding and non-coding RNAs.

This pipeline streamlines the process of quantifying transcript abundance in cells or tissues within their intact anatomical context. Throughput of this pipeline can potentially be increased by using microfluidic devices to array *C. elegans* in ordered locations to speed up imaging (Wan et al. 2019). The high signal-to-noise produced by HCR allows the number of transcript molecules present within a cell to be estimated by counting fluorescent spots. This approach has some standard caveats: the labelling may not be perfectly efficient (Choi et al. 2018), and two closely localized transcripts may not be resolved due to the resolution limits of conventional light microscopy. Super-resolution or expansion microscopy can be used to overcome this latter shortcoming. The detection of transcripts by RS-FISH is deterministic, therefore the only source of variability in the analysis pipeline will be between operator differences in ROI determination. This human error can be minimised by training individuals in robust cell identification.

Despite these caveats, counting transcripts is more accurate and robust than measuring the fluorescence intensities for several technical reasons. Identical transcript molecules may have different fluorescence intensities because they can be labelled with different numbers of fluorophores. Critically, fluorescence levels are sensitive to a multitude of factors in a microscope’s light path from illumination to detection and to operator-dependent variability in ROI boundary delineation. Rigorous fluorescence measurements typically require extensive calibration and normalization. Furthermore, fluorescence measurements are significantly affected by background fluorescence that is prominent in biological tissues. Such background varies tremendously, requires subtraction, is difficult to estimate accurately, and increases the statistical uncertainty of fluorescence estimates due to error propagation in the measurement of two values (signal and background). Moreover, in cells with few transcripts, readouts could be dominated by background, where most pixels in ROI correspond to background. Empirically, our results also show that fluorescence levels are more sensitive to batch effects, which we speculate is due to differences in the number of fluorescent labels attached to the transcript in the amplification step. A key advantage of spot quantification is that it simplifies comparisons, making it is straightforward to compare the abundance of different transcripts in the same cell, or the levels of a given transcript between cells within a sample or between samples. Dispensing with normalization or calibration needed with relative measurements reduces the number of additional measurements needed (thereby reducing workload) and diminishes the uncertainty in quantitative values (since every additional measurement added an associated uncertainty and normalization results in error propagation). While we use background estimates to eliminate false positives, the background values do not contribute directly to the spots counts as they do to every pixel value in fluorescence measurements.

Our results provide examples of how the quantitative transcript counts can be used to reveal differential effects of a mutation on the expression of a given gene among cells as well as heterogeneity among individuals. These results further illustrate the value of our cell-resolved quantitative pipeline ability.

Transcriptional fusions with fluorescent proteins have been widely used in *C. elegans* because of convenience, low costs, and widely available expertise (Boulin et al. 2006). Since estimating RNA abundance is complementary approach, we sought to lower the technical barriers with this quantitative HCR pipeline spanning all the steps from harvesting animals to results comprising transcript counts. The ease of multiplexed experiments provides additional economies of scale. Furthermore, we lower financial barriers by highlighting resources for designing probes and amplifier oligonucleotides that can be bought from third-party suppliers at lower costs. We hope that disseminating this pipeline will increase adoption of quantitative transcript counting in gene expression analysis in *C. elegans* to facilitate more mechanistic studies at the systems level.

## Methods

### *C. elegans* strains and culture

The following strains were used in this study: *N2*, *QL1257 daf-2(e1370)*, *QZ81 ins-6(tm2416)*, and *QZ83 daf-28(tm2308)*. Animals were cultured on standard NGM plates seeded with OP50 *E. coli* food (Stiernagle 2006). For experiments involving L2 larvae, all animals were harvested and fixed within 3 months of thawing from frozen stocks to minimize accumulation of mutations. To control for physiological states and transgenerational effects in the quantitative analysis of *ins-4*, we started with plates of starved arrested animals, grew them with ample OP50 food for exactly 3 generations, then harvested them for HCR analysis. All genotypes were processed simultaneously in parallel to control for batch effects. To collect embryo and adult samples, synchronized L1s were cultured to gravid adults in S-Basal (0.1 M NaCl and 0.05 M potassium phosphate (pH 6.0), 5 µg/ml cholesterol) with E. coli HB101 at a concentration of 1000-3000 worms/mL. To collect embryos, gravid adults from liquid culture were washed twice in M9 buffer and treated with hypochlorite solution (1.2% hypochlorite, 0.5M NaOH) (Stiernagle 2006).

### Fixation and permeabilization

Fixation and permeabilization protocols were adapted from published larval protocols (Choi et al. 2016). Embryos were washed twice in M9 buffer and once in PBS before being resuspended in methanol and stored at -20°C overnight. For fixation, embryos were treated with 50% methanol/PBS, washed in PBST, and fixed in 8% formaldehyde for 10 min at room temperature. This double fixation method prevents nuclear collapse (Supplemental Figure 1). After washing, samples were incubated for 15 min in glycine (2 mg/mL) on ice and washed again before proceeding to hybridisation.

L2 larvae were washed three times in M9 buffer and fixed in 4% formaldehyde followed by storage at −80°C overnight. Following thawing, larvae were washed in PBST, treated with proteinase K (100μg/mL, 10 min at 37°C), washed again, and incubated in glycine (2 mg/mL) on ice. Samples were washed again before proceeding to hybridisation.

Adult worms were washed in PBS, kept in methanol at -20°C overnight, incubated in 50% methanol/PBS for 5 in at room temperature before fixation. Samples were fixed in 8% formaldehyde, washed, treated with glycine, and permeabilised with proteinase K (100μg/mL, 10 min at 37°C). Following additional washes, worms were post-fixed in 70% ethanol at 4°C overnight and used within one week.

### HCR probe design, hybridization and staining

Probes were designed and ordered as custom oligopools (Integrated DNA Technologies), resuspended in DEPC-ddH₂O at 1μM concentration, incubated at 4°C overnight, and stored at - 20°C. Fluorophore-labelled HCR hairpin amplifiers (h1 and h2) corresponding to specific initiator sequences (B isoforms) were designed based on published sequences (Choi et al. 2014) or ordered from Molecular Instruments. Hairpins were resuspended to 3μM concentration in hairpin resuspension buffer, aliquoted, and stored at -20°C.

Samples were washed in PBST and equilibrated in probe hybridisation buffer. Pre-hybridisation was performed in probe hybridisation buffer at 37°C for 1h. Probe solutions were prepared by adding 2pmol of each probe pair mix to hybridisation buffer. Samples were incubated with probes (final volume 500μL) overnight (>12 h) at 37°C. The next day probes were removed by 4 x 15 min washes in pre-warmed probe wash buffer at 37°C, followed by two washes in 5× SSCT at room temperature.

Samples were equilibrated in amplification buffer for 30 min at room temperature. Hairpin amplifiers (h1 and h2) were snap-cooled (95°C for 90 s, then cooled to room temperature in the dark for 30 min) prior to use.

Hairpin solutions were prepared by combining equal volumes of h1 and h2 for each fluorophore channel and diluting in amplification buffer. Samples were incubated with hairpin solution (final volume 500μL) overnight (>12 h) at room temperature in the dark.

Excess hairpins were removed by sequential washes in 5× SSCT, before staining with DAPI (1μg/mL in 5× SSCT) for 1h at room temperature in the dark, followed by a final 5× SSCT wash. Samples stored in mounting medium and stored at 4°C protected from light prior to imaging (full protocol in supplementary).

The L2 larval protocol produced positive staining in most animals across 3 independent experimental batches (∼90%; Supplemental figure 2).

### Microscopy

Larval images were acquired with on Zeiss LSM 800 confocal microscope with a x63 1.4NA oil immersion objective with wavelengths 405 (lasers at 0.15%, gain at 650v), 488 (lasers at 2%, gain at 650v), 561 (lasers at 3%, gain at 700v) and 640 nm (lasers at 2%, gain at 675v). Dwell time was 1.03μs, with a 0.5 μm z-distance and line averaging at 2x. Spectral separation was achieved exploiting differences in wavelengths.

Embryo and adult images were acquired on a Nikon AX R confocal microscope equipped with a resonant scanner and 60x 1.4NA oil immersion objective with wavelengths 488, 546, and 647 nm lasers at 3%, with gain at 30v and line averaging at 16x.

Fluorescence imaging was performed using a 405nm wavelength for DAPI in all samples. The following fluorophores were used for each developmental stage: **embryo** *ref-1* (Alexa Fluor 488), *pes-10* (Alexa Fluor 546) and *end-1* (Alexa Fluor 647); **larvae** *ins-6* (Alexa Fluor 488), *ins-4* (Alexa Fluor 546) and *daf-28* (Alexa Fluor 647); **adult** *nos-1* (Alexa Fluor 546).

### Image Analysis

For image analysis in larvae, a 3D image stack was taken with the focal planes imaged determined using DAPI to find the dorsal and ventral points of the worm, ensuring the entire worm was captured. RS-FISH (Bahry et al. 2022) was applied to identify fluorescent spots at single-molecule resolution. Prior to automated detection, the volumes of the worm were delineated using binary masks generated from Alexa Flour 488 levels via automated thresholding, morphological operations, and selection of the largest connected region. Parameters including the anisotropy coefficient and spot size were manually optimized for each imaging channel. Spot intensity thresholds were dynamically set for each image as the mean signal intensity plus three standard deviations, calculated within the masked region. RANSAC fitting was used for spot detection. Local background subtraction was applied using RANSAC on mean. A customized ImageJ macro script was used to execute the above steps in batch mode.

To assess whether detected objects remained spatially resolvable following overnight HCR amplification we measured their full width at half maximum (FWHM) along the X/Y dimension in a random sample (Supplemental Figure 5). The average lateral FWHM was ∼ 0.3 μm and >90% of objects had a lateral FWHM below 0.5 μm. These values are close to the practical resolution of a 1.4 NA objective, indicating most objects remain diffraction limited following overnight signal amplification.

Owing to the complex cellular morphology, cell segmentation was performed manually. Individual cells were delineated based on their boundaries in the XY maximum projection and their extent along the Z-axis. Cell boundaries were traced using the freehand selection tool in FIJI (Schindelin et al. 2012), and the corresponding XY coordinates were exported. Cell identities and Z-plane positions were subsequently recorded in an Excel spreadsheet.

To calculate cell properties, a custom MATLAB script was developed by Dr. Jane Ling and proofread by Claude (Anthropic, accessed 2026) for code simplification. The script employed a convex hull algorithm to determine whether each detected spot was located within a defined cell boundary. To assess spot-specific signal, spot coordinates were mapped onto a voxel grid spanning the cell volume to generate a binary mask, which was subsequently dilated using an anisotropic structuring element reflecting expected spot dimensions. Intensities within this dilated mask were used to calculate total puncta intensity, while the remaining voxels within the cell were treated as local background. The total number of fluorescent spots and associated intensity metrics were then quantified per cell and summarized in the output table.

Three-dimensional interactive visualization of the segmented cell boundaries and multi-channel fluorescence microscopy data was performed using Napari (Sofroniew et al. 2026), an open-source image viewer built on Python. This interactive visualization platform enabled rapid assessment of segmentation accuracy and inspection of fluorescence intensity distributions within individual cells across the three-dimensional volume.

For all quantitative analysis we only included data on days of imaging where the *ins-4* channel (Alexa Fluor 546) was not saturated i.e. If any animal on a given day had saturation in the *ins-4* channel, the entire data from that day was discarded. Saturation was classified as the center point of any *ins-4* spot within the sample having a fluorescence value of 250 a.u. or above.

### Statistical Analysis

All statistical analyses were performed in R (Team 2024) using the lme4 (Bates et al. 2015) and tidyverse (Wickham et al. 2019) packages. The codes were written by Dr. Fursham Hamid, then proofread with Claude (Anthropic, accessed 2026) before being reviewed manually by Dr. Hamid and tested on dummy data sets. Significant difference in corrected mean fluorescence and transcript counts of each ILP mRNA between genotypes was determined using a generalized linear model (GLM). Continuous (corrected mean fluorescence) and integer (transcript counts) quantitative data were fitted to a Gamma or Negative Binomial distributions respectively due to the high variance nature of the dataset. For comparisons that involve animals taken from distinct batches, a mixed-effect model (GLMM) was used that included the experimental batch as a random effect variable. We analysed data from individual neurons but chose to focus on analysing ASI and ASJ data in their neuron pairs, including the cell side (L/R) in the mixed effect model as a fixed effect. Model-derived estimates were used to calculate mean expression levels for mutant and control groups and log₂ fold changes (log2FC) were computed from model coefficients. Statistical significance was assessed using z-statistics, and corresponding p-values were reported for genotype effects, after Benjamini–Hochberg correction for multiple testing.

We examined how much of the model-scale variance for corrected mean fluorescence and total spot count could be assigned to experimental batch, neuron side, fixed effects and unexplained variation (Supplemental Table 1). Raw variance estimates cannot be compared between the two models as they were analysed using different distributions and transformations. Therefore, we compared the proportion of model-scale variance that could be attributed to each component. In the same samples the corrected mean fluorescence model had experimental batch account for 28.8% of the variance, whereas it only accounted 2.2% of the variance in the transcript count mode, showing substantially less between batch variation.

## Supporting information

Protocol

## Code availability

The Image analysis pipeline and codes are published on GitHub https://github.com/f-hamidlab/HCR_worm.

## Acknowledgements

We acknowledge the use of Artificial Intelligence in aspects of the data analysis as detailed in the methods.

We thank the following people at the Centre for Developmental Neuroscience at King’s College London: Corinne Houart and Joshua Bradbury for help with HCR probe design and set up, Darren Williams and Connor Sproston for help with HCR hairpins, and Stephen Ogg for microscopy support. We would like to thank Chintan Trivedi at University College London Imaging Facility for allowing us to use his probe design tool, Nicola Lawrence at the Gurdon Institute Imaging Facility at Cambridge University for microscopy support, and Joy Alcedo for *C. elegans* strains. We are grateful to Hang Lu for comments on the manuscript. JA acknowledges funding from the Wellcome Trust (217170). QC acknowledges funding from the Leverhulme Trust (17680 & 23722).

---

**Supplemental Figure 1.**
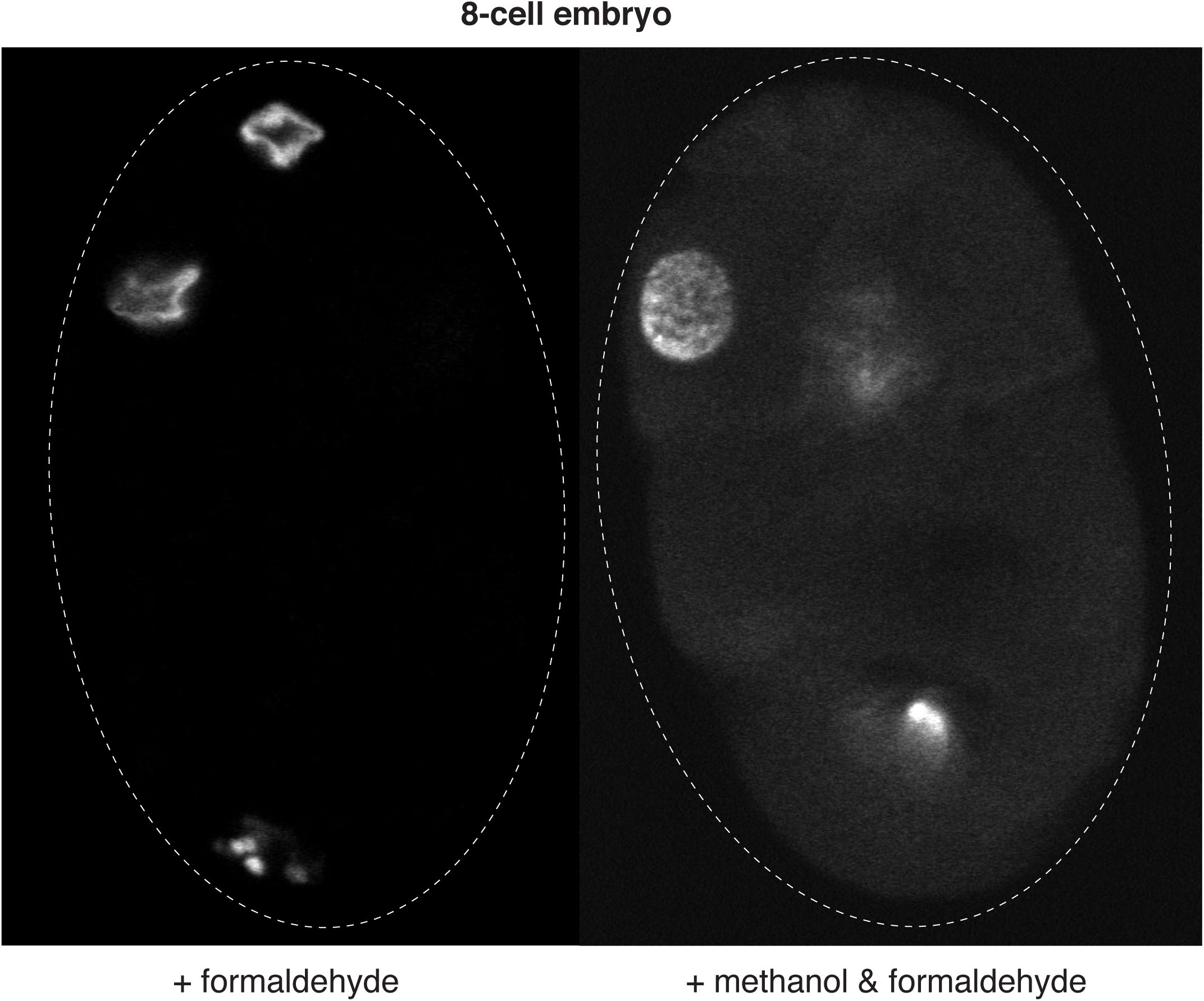
Nuclear morphology in different fixation methods. Representative images of 8-cell stage embryos fixed with formaldehyde alone (left) or methanol and formaldehyde (right). Nuclei (DAPI, grey) collapse following formaldehyde treatment, whereas nuclear morphology is better preserved with formaldehyde and methanol treatment. Selected cells (C and P3) are indicated. Dashed outlines mark embryo boundaries.

**Supplemental Figure 2.**
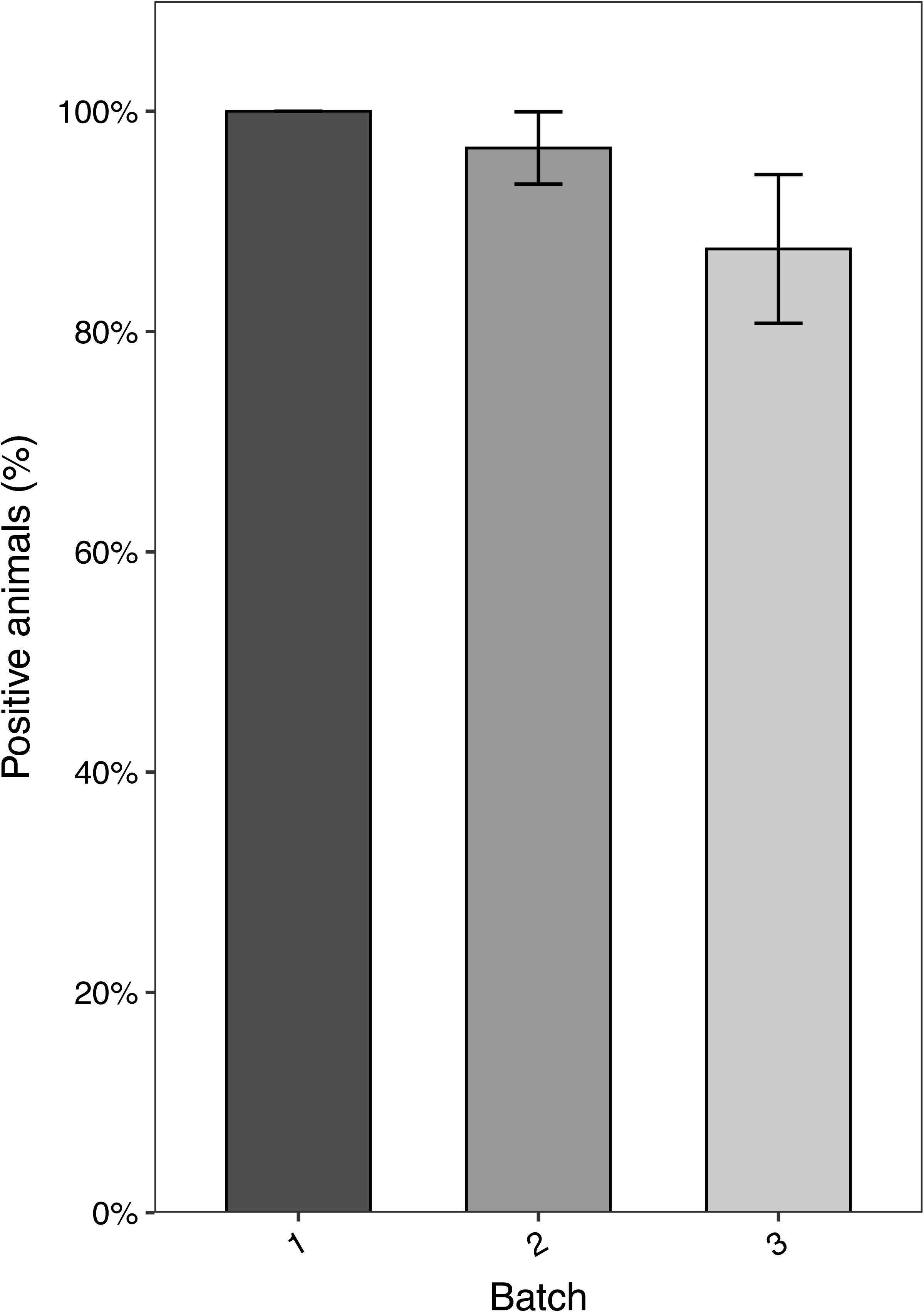
Staining efficiency in L2 protocol. Percentage of L2 wild type larvae exhibiting positive signal following fixation and HCR3.0 staining across 3 experimental batches. A larva was considered positively stained if there was staining in all 3 imaging channels.

**Supplemental Figure 3.**
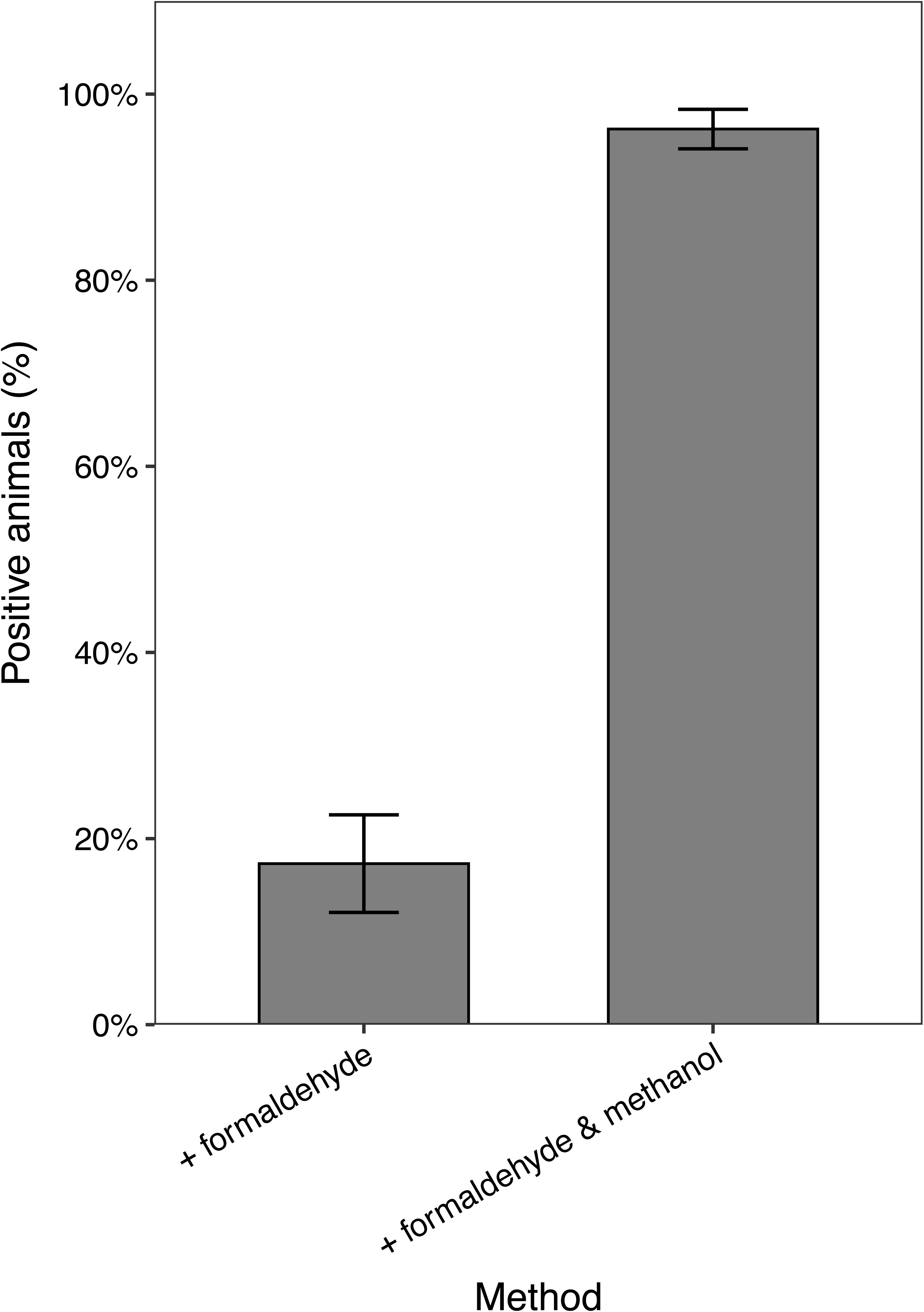
Comparison of fixation methods for signal detection. Percentage of adults exhibiting positive signal following fixation with formaldehyde alone or with methanol and formaldehyde.

**Supplemental Figure 4.**
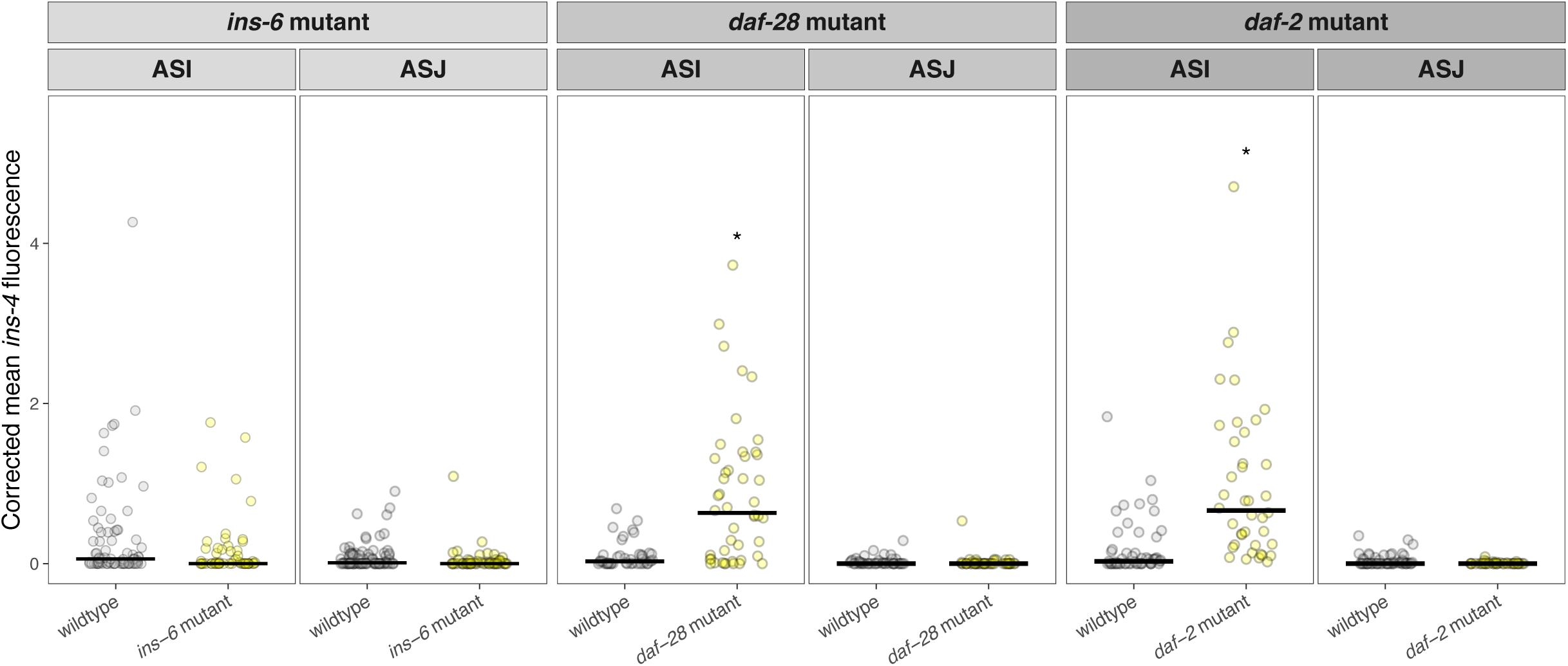
Corrected mean fluorescence quantification. *ins-4* fluorescence quantification in individual ASI and ASJ neurons from wildtype and mutant L2 larvae. Columns show the mutant background: *ins-6(tm2416)*, *daf-28(tm2308)* and *daf-2(e1370)*. Grey and yellow points indicate wildtype and mutant genotype respectively. Each point represents the transcript count from a left and right neuron pair; horizontal black bars indicate the median. Asterisks indicate statistically significant differences between wildtype and mutant animals (*P < 0.05).

**Supplemental Figure 5.**
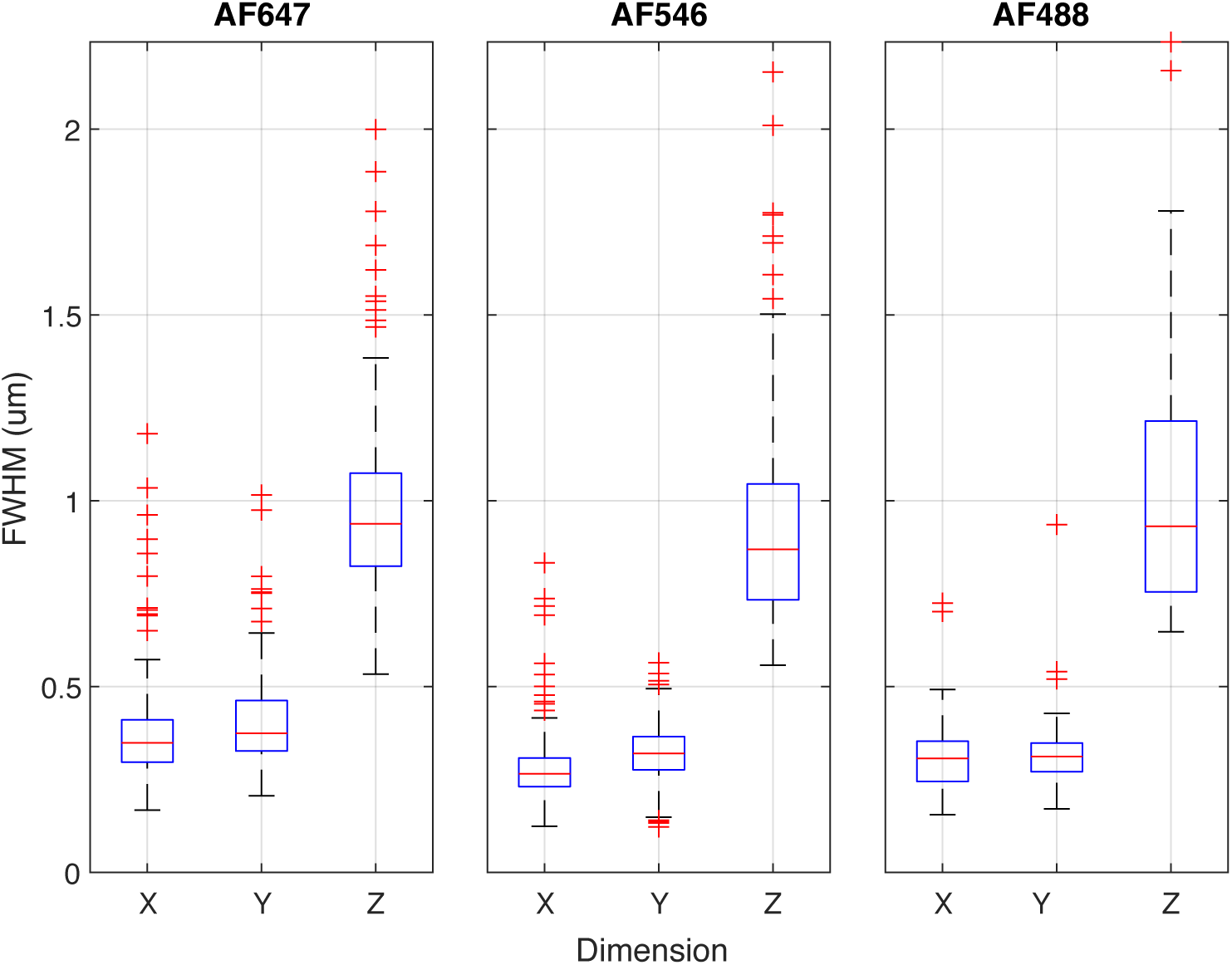
Dimensions of RS-FISH detected objects. Distribution of the full width at half maximum (FWHM, µm) of detected objects across the X, Y, and Z dimensions in three fluorescence channels (AF647, AF546, AF488). Boxes show the interquartile range, horizontal red lines indicate the median, and red whiskers show values within 1.5 times the interquartile range.

**Supplemental Table 1.** Variance across models.

| Model | Cell type | Position % | Batch variance | Batch % | Fixed effect variance | Unexplained variance | Total model scale variance |
| --- | --- | --- | --- | --- | --- | --- | --- |
| Transcript count | 0.00033 | 0.02% | 0.04299 | 2.21% | 0.640997 | 1.260723 | 1.94471 |
| Total intensity of spots | 0.00444 | 1.36% | 0.03248 | 9.95% | 0.122784 | 0.171165 | 0.32643 |
| Corrected mean fluorescence | 0 | 0% | 1.09485 | 28.77% | 0.997465 | 1.712924 | 3.805243 |

**Supplemental Table 2.** *ins-4* transcript count quantification summary.

| Comparison | Cell type | Genotype | n | number of 0 values | median | BH-adjusted p value |
| --- | --- | --- | --- | --- | --- | --- |
| *ins-6* mutant | ASI | wildtype<br>mutant | 76<br>54 | 26<br>28 | 2<br>0 | 0.1721 |
|  | ASJ | wildtype<br>mutant | 76<br>54 | 31<br>33 | 1<br>0 | 0.0006347 |
| *daf-28* mutant | ASI | wildtype<br>mutant | 42<br>46 | 15<br>6 | 1<br>17 | 1.97E-08 |
|  | ASJ | wildtype<br>mutant | 42<br>46 | 23<br>34 | 0<br>0 | 0.1721 |
| *daf-2* mutant | ASI | wildtype<br>mutant | 54<br>40 | 22<br>0 | 1.5<br>19 | 1.97E-08 |
|  | ASJ | wildtype<br>mutant | 54<br>40 | 29<br>30 | 0<br>0 | 0.0005525 |

## Notes

https://github.com/f-hamidlab/HCR_worm

### Competing Interest Statement

The authors have declared no competing interest.

### Summary of Updates

Results and Figures 3 and 5 now focus on ins-4 expression rather than ins-6. Comparisons of transcript counting and fluorescence intensity measurements for ins-4 have been added, along with analysis of batch effects. Supplementary figures and tables have been added to document staining efficiency, spot dimensions and fluorescence intensity quantitative results. The Introduction, Methods, Results, Discussion and figure legends have been revised to reflect the changes listed above.

