## Supplementary material for "An integrated pipeline to estimate transcripts counts with single-cell resolution in *Caenorhabditis elegans*": Protocol

***C. elegans* RNA Hybridisation Chain Reaction (HCR) Protocol**

**Solutions and Buffers**

DEPC ddH_2_O 1L

1L ddH_2_O

Add 1ml DEPC and leave benchtop overnight for >12 hours then autoclave.

1M MgSO_4_ DEPC 50ml

12.34g MgSO_4_

Fill up to 50ml with DEPC ddH_2_O and autoclave.

1M Tris DEPC 100mL

12.114g Tris

HCL to pH 7.0

Fill up to 100ml with DEPC ddH_2_O and autoclave.

0.5M EDTA DEPC 100mL

18.53g EDTA

NaOH to pH 8.0

Fill up to 100ml with DEPC ddH_2_O and autoclave.

3M NaCl DEPC 100mL

17.53g NaCl

Fill up to 100ml with DEPC ddH_2_O and autoclave.

Hairpin Resuspension Buffer 40mL:

400μL 1M Tris DEPC

80μL 0.5M EDTA DEPC

4mL 3M NaCl DEPC

Fill up to 50ml with DEPC ddH_2_O.

M9 DEPC 1L

3g KH_2_PO_4_

6g Na_2_HPO_4_

5g NaCl

1mL 1M MgSO_4_ DEPC

Fill up to 1L with DEPC ddH_2_O and autoclave.

Store at 4°C.

4% Paraformaldehyde (PFA) DEPC 40ml

10ml 16% PFA solution

4ml 10x PBS

Fill up to 40ml with DEPC ddH_2_O, aliquot and store at -20°C.

PBST DEPC 50ml

5ml 10x PBS

500μL 10% Tween 20

Fill up to 50ml with DEPC ddH_2_O

1x PBS DEPC/ 50%Methanol 3mL

1.5mL Methanol

300μL 10X PBS

1.5mL DEPC ddH_2_O

Proteinase K (100 μg/mL) Solution 1mL

5μL of 20 mg/mL proteinase K

995μL PBST DEPC

Glycine Solution 50ml

100mg Glycine

Fill up to 50ml with PBST DEPC

5xSSCT DEPC 50ml

12.5ml 20x SSC

500μL 10% Tween 20

Fill up to 50ml with PBST DEPC

5x SSCT DAPI DEPC 1mL

1 μL 1mg/mL DAPI stock

995μL 5xSSCT DEPC

2 x Fixation buffer (8% Formaldehyde) 10mL

5mL 16% Formaldehyde

1mL 10X PBS

4mL DEPC ddH_2_O

Probe pairs (P1 and P2) for each gene of interest

Specific probes complementary for a given target RNA sequence are generated using a custom HCR probe designer (Suppermpool et al, 2025). P1 and P2 carry half of the HCR initiator sequence, and when both pairs bind to the target RNA the full initiator sequence is exposed. When designing probes, you select for specific imitator sequences that match to specific fluorophore b isoforms. It is recommended to have >15 probe pairs, however we have successful HCR with as few as 4 probe pairs. The probe pairs are ordered from IDT in a custom oligopool at a concentration of 50pmol. The oligo pool is resuspended in 50μL of DEPC ddH_2_O and left at 4°C for >12 hours before storing at -20°C.

Hairpin Amplifiers (h1 and h2) 3μM stocks

HCR uses pairs of reporter-labelled hairpin amplifiers that bind to the exposed HCR initiator sequence, triggering a self-amplification cascade and creating a reporter-labelled amplification polymer attached to the target RNA molecule. These hairpins come in different B isoforms that correspond to specific fluorophore wavelengths. The fluorophore and B isoform are selected during probe design based on the wavelength availability in imaging set ups. HCR3.0 hairpins can be ordered ready to use from Molecular Instruments or ordered from IDT as dual HPLC purified lyophilised custom oligos using published sequences for the B isoforms (Choi et al, 2014) with alexa fluorophores on the end of the 3’ and 5’ sequence. Resuspended a 3uM concentration in Hairpin Resuspension Buffer, aliquoted and stored at -20°C.

The following buffers are supplied by Molecular instruments HCR3.0 kit (NOT the new HCR Gold RNA-FISH kits as probes and hairpins not compatible):

- Probe Hybridization Buffer
- Probe Wash Buffer
- Amplification Buffer

These can also be made in house using the following protocols (Choi et al, 2014):

Probe hybridisation buffer 40ml

20mL formamide

10mL 20x SSC

360μL 1M citric acid, pH 6.0

400μL of 10% Tween 20

200μL of 10mg/mL heparin

800μL of 50% dextran sulfate

Fill up to 40ml with PBST DEPC

Probe wash buffer 40ml

20mL formamide

10mL 20x SSC

360μL 1M citric acid, pH 6.0

400μL of 10% Tween 20

200μL of 10mg/mL heparin

Fill up to 40ml with PBST DEPC

Amplification buffer 40ml

10mL 20x SSC

400μL of 10% Tween 20

8mL of 50% dextran sulfate

Fill up to 40ml with PBST DEPC

Antifade Mounting Medium

We use EverBrite™ Mounting Medium with DAPI from Biotum (23002) or Vectashield Antifade Mounting medium with DAPI from Vector Laboratories (H-1200).

**Protocol**

Collection and fixation and probe hybridisation steps are stage dependent as follows:

**(A) Embryos**

- Embryo centrifugation is performed at 800g for 1 min, except when the embryos are in viscous Amplification or Hybridisation buffer, which require 1800g for 1 min.

**Harvesting animals**

Day 1

1. Perform experiment-specific treatments and collect embryos by hypochlorite treatment to obtain samples of ~1000-2000 embryos.
2. Wash 2 times with 1mLof M9 DEPC buffer. Centrifuge at 800g/0.8rcf for 1 min to pellet embryos between washes.
3. Resuspend in 1mL ice-cold methanol.
4. Place in -20°C overnight (>12hr) before use. Frozen embryos can be stored long-term at -20°C.

**Fixing and permeabilisation**

Day 2

5. Pellet embryos at 800g/0.8rcf and remove methanol.
6. Resuspend in 1ml 1X PBS DEPC/50% methanol, incubate for 5min at room temperature.
7. Pellet embryos and remove 1xPBS DEPC/50% methanol. Resuspend in 1ml of PBST DEPC and incubate at room temperature for 5 min.
8. Wash the embryos with 1mL of PBST DEPC. Centrifuge at 800g/0.8rcf for 1 min to pellet then remove PBST DEPC.
9. Add 500μL of PBST DEPC and 500μL of 2X fixation buffer to each tube and mix well.
10. Incubate for 10min at room temperature
11. Wash 2 times with 1mL of PBST DEPC. Centrifuge at 800g/0.8rcf for 1 min between washes to pellet and after remove PBST DEPC.
12. Incubate in 1mL of ice-cold 2 mg/mL Glycine Solution for 15min on ice.
13. Wash 2 times with 1mL of PBST DEPC.
14. Continue to probe hybridisation step.

**(B) L2 stage larvae**

**Harvesting animals**

Day 1

1. Perform experiment-specific treatments to obtain L2 larvae samples. We aim to harvest ~500 L2 larvae per sample.
2. Wash L2 Larvae off plate(s) with appropriate amount of M9 DEPC buffer into to 2 x 1.5mL eppendorfs. For 5 small (6cm) plates we use 2mL of M9 DEPC buffer.
3. Wash each tube of larvae 3 times with 500μL of M9 DEPC buffer. Centrifuge at 200g/0.2rcf for 2min to pellet larvae between washes.
4. Centrifuge and remove ~400μL of M9 buffer.
5. Add 500μL of 4% paraformaldehyde (PFA).
6. Place in -80°C overnight (>12hr) before use. Frozen worms can be stored long-term at -80°C.

**Fixing and permeabilisation**

Day 2

7. Fix larvae by thawing at room temperature for 45min.
8. Wash 2 times with 500μL of PBST DEPC. Centrifuge at 200g/0.2rcf for 2min between washes to pellet larvae.
9. Treat larvae with 1mL of Proteinase K Solution for 10min at 37°C.
10. Wash larvae 2 times with 500μL of PBST DEPC. Centrifuge at 200g/0.2rcf for 2min between washes to pellet larvae.
11. Incubate in 1mL of 2 mg/mL Glycine Solution for 15min on ice.
12. Pre-warm Probe Hybridization Buffer: 500μL at 37°C and 250μL at room temperature per sample for subsequent probe hybridisation steps.
13. Wash larvae 2 times with 500μL of PBST DEPC. Centrifuge at 200g/0.2rcf for 2min between washes to pellet larvae.
14. Continue to probe hybridisation step.

**(C) Adults**

- Adult centrifugation is performed at 375g for 2 minutes.

**Harvesting animals**

Day 1

1. Perform experiment-specific treatments to obtain adult samples. We aim to harvest ~500 adults per sample.
2. Wash adults off plate(s) with appropriate amount of M9 DEPC buffer into to a flacon tube.
3. Wash the adults 2 times with 1mL of M9 DEPC. Centrifuge at 375g/0.4rcf for 2min to pellet adults between washes.
4. Centrifuge and remove M9 DEPC.
5. Resuspend in 1mL of methanol.
6. Place in -20°C overnight (>12hr) before use. Frozen worms can be stored long-term at -20°C.

**Fixing and permeabilisation**

Day 2

7. Pellet adults at 375g/0.4rcf and remove methanol.
8. Resuspend in 1ml 1X PBS DEPC/50% methanol, incubate for 5min at room temperature.
9. Pellet embryos and remove 1xPBS DEPC/50% methanol. Resuspend in 1ml of PBST DEPC and incubate at room temperature for 5 min.
10. Wash the adults with 1mL of PBST DEPC. Centrifuge at 375g/0.4rcf for 2min to pellet then aspirate PBST DEPC.
11. Add 500μL of PBST DEPC and 500μL of 2X fixation buffer to each tube and mix well.
12. Incubate for 10min at room temperature
13. Wash 2 times with 1mL of PBST DEPC.
14. Incubate in 1mL of ice-cold 2 mg/mL Glycine Solution for 15min on ice.
15. Wash 2 times with 1mL of PBST DEPC.
16. Resuspend in 995μL of PBST DEPC and add 5μL 20 mg/mL proteinase K. Incubate for 10min at 37°C.
17. Wash 2 times with 1mL of PBST DEPC.
18. Remove PBST DEPC and resuspend in 1mL of 70% ethanol. Leave at 4°C overnight. Fixed adult worms can be stored at 4°C and used for up to 1 week.
19. Was 2 times with 1ml of PBST DEPC.
20. Continue to probe hybridisation step.

HCR Probe hybridisation and amplification steps are standardised independent of stage:

- Embryo centrifugation is performed at 800g, except when the embryos are in viscous Amplification or Hybridisation buffer, which require 1800g.
- Larval centrifugation is performed at 500g/0.5rcf.
- Adult centrifugation is performed at 375g for 1 min.

**Probe hybridisation**

Day 2 (embryo and larvae)/ 3 (adult)

Pre-warm probe hybridization buffer to 37C

1. Incubate sample in 1 mL of 50% PBST / 50% probe hybridization buffer for 5 min at room temperature.
2. Centrifuge and aspirate buffer.
3. Pre-hybridize samples in 300μL of Probe Hybridization Buffer at 37°C for 1hr.
4. Prepare Probe Solution by adding 2pmol (2.1μL of a 1μM stock) of each probe to 210μL of Probe Hybridization Buffer at 37°C.
5. Add 200μL Probe Solution to the sample to reach a final hybridization volume of 500μL.
6. Incubate overnight (>12hr) at 37°C.

Day 3 (embryo and larvae)/ 4 (adult)

1. Pre-warm Probe Wash Buffer 15min before washes at 37°C.
2. Add 1 mL probe wash buffer, invert to mix. For embryos, spin down at 3000g at this step. The usual spin for adults and larvae.
3. Remove excess probes by washing 4 x 15min with 1mL pre heated probe wash buffer at 37°C. Centrifuge between washes.
4. Pre-warm probe amplification buffer: 500μL at room temperature per sample.
5. Wash 2 times for 5min with 1mL 5xSSCT DEPC at room temperature. Centrifuge for 2min between washes.
6. Pre-amplify samples with 300μL of Amplification Buffer for 30min at room temperature.
7. Thaw all Hairpin Amplifier for each B isoform in use: each B isoform has two Hairpin Amplifiers (h1 and h2). Add 10μL of the first amplifier hairpin h1, per sample, in a sterile PCR tube. In a separate tube add an equal volume of amplifier hairpin h2. Repeat this step for each distinct B isoform that was used during probe hybridization. Carefully label each tube with the corresponding hairpin number and B isoform.
8. Snap-cool the amplifier hairpins by heating all Hairpin Amplifier tubes at 95°C for 90s and then let them cool to room temperature in the dark for 30min.
9. Prepare Hairpin Solution by adding snap-cooled B isoform h1 and h2 hairpins to Amplification Buffer isoform at room temperature to a final volume of 200µL (e.g. If you have 3 x B isoforms per sample you will be mixing 60 µL of cooled hairpins with 140µL of Amplification Buffer).
10. Add the Hairpin Solution to sample to reach a final volume of 500μL.
11. Incubate the samples in the dark at room temperature by wrapping with aluminium foil (3 hours for embryos, overnight (>12hr) for larvae and adults).

HCR Probe washes and amplification

Day 4 (embryo and larvae)/ 5 (adult)

12. Add 1 mL 5xSSCT DEPC, invert to mix. For embryos, spin down at 3000g at this step. The usual spin for adults and larvae.
13. Remove excess hairpins by washing with 1mL 5x SSCT DEPC in the dark at room temperature:

(a) 2 times for 5 min

(b) 2 times for 30min

14. Freshly prepare 1mL DAPI in 5xSSCT DEPC at a concentration of 1μg/mL (1μL of 1mg/mL DAPI stock in 1mL 5xSSCT DEPC).
15. Add the DAPI solution to the larvae and incubate for 30 min - 1hr in the dark at room temperature.
16. Do a final wash of the larvae for 5min with 1 ml of 5xSSCT DEPC in the dark at room temperature.
17. Centrifuge and remove 5xSSCT DEPC.
18. Add desired volume of preferred Mounting Medium and store at 4°C protected from light by wrapping in aluminium foil before microscopy.
